# Poly(A) binding protein is required for mRNP remodeling to form P-bodies in mammalian cells

**DOI:** 10.1101/2021.02.07.430159

**Authors:** Jingwei Xie, Yu Chen, Xiaoyu Wei, Guennadi Kozlov

## Abstract

Compartmentalization of mRNA through formation of RNA granules is involved in many cellular processes, yet it is not well understood. mRNP complexes undergo dramatic changes in protein compositions, reflected by markers of P-bodies and stress granules. Here, we show that PABPC1, albeit absent in P-bodies, plays important role in P-body formation. Depletion of PABPC1 decreases P-body population in unstressed cells. Upon stress in PABPC1 depleted cells, individual P-bodies fail to form and instead P-body proteins assemble on PABPC1-containing stress granules. We hypothesize that mRNP recruit proteins via PABPC1 to assemble P-bodies, before PABPC1 is displaced from mRNP. Further, we demonstrate that GW182 can mediate P-body assembly. These findings help us understand the early stages of mRNP remodeling and P-body formation.

**Summary statement:** A novel role of poly(A) binding protein is reported in P-body formation

## Introduction

Cytoplasmic messenger ribonucleoproteins (mRNPs) are suggested to cycle among polysomes, stress granules (SGs) and P-bodies (PBs). In this cycle, mRNAs exist in different functional states from translating, non-translating to degradation. While polysomes consist of translating mRNAs and SGs paused mRNAs with translation initiation components, PBs contain mRNA decay machineries.

PBs are present in unstressed cells, and further induced upon inhibition of translation initiation (Teixeira et al., 2005, Kedersha et al., 2005). PBs are closely related to control of translation and mRNA degradation. Proteins at PBs are involved in mRNA decay and translation repression, including decapping enzyme complex Dcp1/Dcp2; decapping activator hedls/GE-1; translation repressor and decapping activator DDX6/RCK, and the CCR4/NOT deadenylase complex (Anderson and Kedersha, 2006, Eulalio et al., 2007a).

SGs can be juxtaposed with PBs in animal cells(Kedersha et al., 2005, Stoecklin and Kedersha, 2013), which suggests that they are close in origin. SGs share some common components with PBs, but distinctively contain translation initiation factors like PABPC1, eIF4G, eIF4A, eIF3 and eIF2 etc. (Decker and Parker, 2012).

Assembly of PBs and SGs may start from non-translating mRNPs, which aggregate into microscopic granules through certain protein-protein interactions. In yeast, a self-interacting domain of Edc3 protein and prion-like glutamine/asparagine (Q/N) rich region of Lsm4 can facilitate the aggregation (Reijns et al., 2008, Decker et al., 2007). However, in metazoans, Edc3 is not required for PB assembly (Eulalio et al., 2007b). Instead, depletion of GW182 or hedls/GE-1, two proteins containing low-complexity and Q/N rich regions, leads to decreased PBs in unstressed animal cells (Eulalio et al., 2007b, Liu et al., 2005a, Yu et al., 2005, Kato et al., 2012).

One interesting issue is the absence of most translation initiation factors, including poly(A) binding protein cytoplasmic 1 (PABPC1), in P-bodies of mammalian cells. Components of RNA granules are in dynamic exchange with cytoplasmic proteins (Kedersha et al., 2005, Andrei et al., 2005). The association and dissociation of proteins, such as PABPC1, to and from the cap and tail of mRNA may be an important step in the transitions of mRNAs from translating to non-translating or decay states.

mRNAs have a poly(A) tail of about 50-100 nucleotides in mammalian cells (Chang et al., 2014, Subtelny et al., 2014). PABPC1 occupies around 27-residues on the poly(A) tail and forms repeating structures (Baer and Kornberg, 1983). PABPC1 is the major isoform out of four known in mammals. PABPC1 consists of four RNA binding domains (RRM 1-4) followed by a poorly conserved linker region and a protein-protein interaction (MLLE) domain at the C-terminus. The RRM domains are pivotal for circularization of mRNA through the binding of the poly(A) tail and eIF4G (Kahvejian et al., 2005, Deo et al., 1999, Imataka et al., 1998, Safaee et al., 2012). The MLLE domain recognizes a conserved PAM2 peptide motif, found in a number of proteins including PB components GW182, Pan3 and Tob1/2 (Xie et al., 2014).

PABPC1 plays important roles in decay besides translation of mRNA. Yeast poly(A) binding protein (Pab1) can inhibit Ccr4/Pop2/Not deadenylase complex (Tucker et al., 2002), possibly due to increased association and protection of Pab1 to the poly(A) tail. It was recently shown that PABPC1 helps recruiting microRNA-induced silencing complex (miRISC) through GW182, a major component of miRISC (Moretti et al., 2012). GW182, in turn, facilitates PABPC1 dissociation from silenced mRNA without deadenylation (Zekri et al., 2013).

Although PABPC1 displacement is a critical step in mRNA decay and can act as a scaffold protein on mRNA to recruit PB proteins GW182 and Pan3 etc. through PAM2 motifs, it is not known what role PABPC1 plays in assembling PBs. Here, we report that depletion of PABPC1 affects PB numbers in unstressed cells and causes fusion of PB and SG components upon cell stress. Further, we found that the availability of PABPC1, but not the presence of SG structures, is critical in PB genesis. By engineering GW182 to strengthen its binding to PABPC1, we simulated the fusion of SGs and PBs. Overall, we conclude that PABPC1 serves as a platform for early mRNP remodeling to form PBs.

## Results

### PABPC1 protein depletion decreases PBs in unstressed cells

We were able to knock-down PABPC1 using siRNAs as previously shown (Yoshida et al., 2006). The number of PBs decreased significantly upon PABPC1 knock-down in unstressed HeLa and MEF cells (Fig. 1A). PABPC1 depletion affects bulk translation only mildly and doesn’t perturb the levels of the major translation factors (Yoshida et al., 2006). However, Paip2, a PABPC1-interacting protein, decreases in cells following PABPC1 depletion (Yoshida et al., 2006). To exclude a role of Paip2 in PB formation, we knocked down both isoforms, Paip2a and Paip2b, by siRNA and found little effect on the number of PBs (Fig. 1B). It was reported that depletion of hedls or GW182 decreased PBs in cells (Eulalio et al., 2007b, Liu et al., 2005b, Yu et al., 2005). However, PABPC1 depletion didn’t lower hedls or GW182 protein levels (Fig. 1C), which suggests that PABPC1 is affecting PBs independently of changes in hedls or GW182.

**Figure 1.**
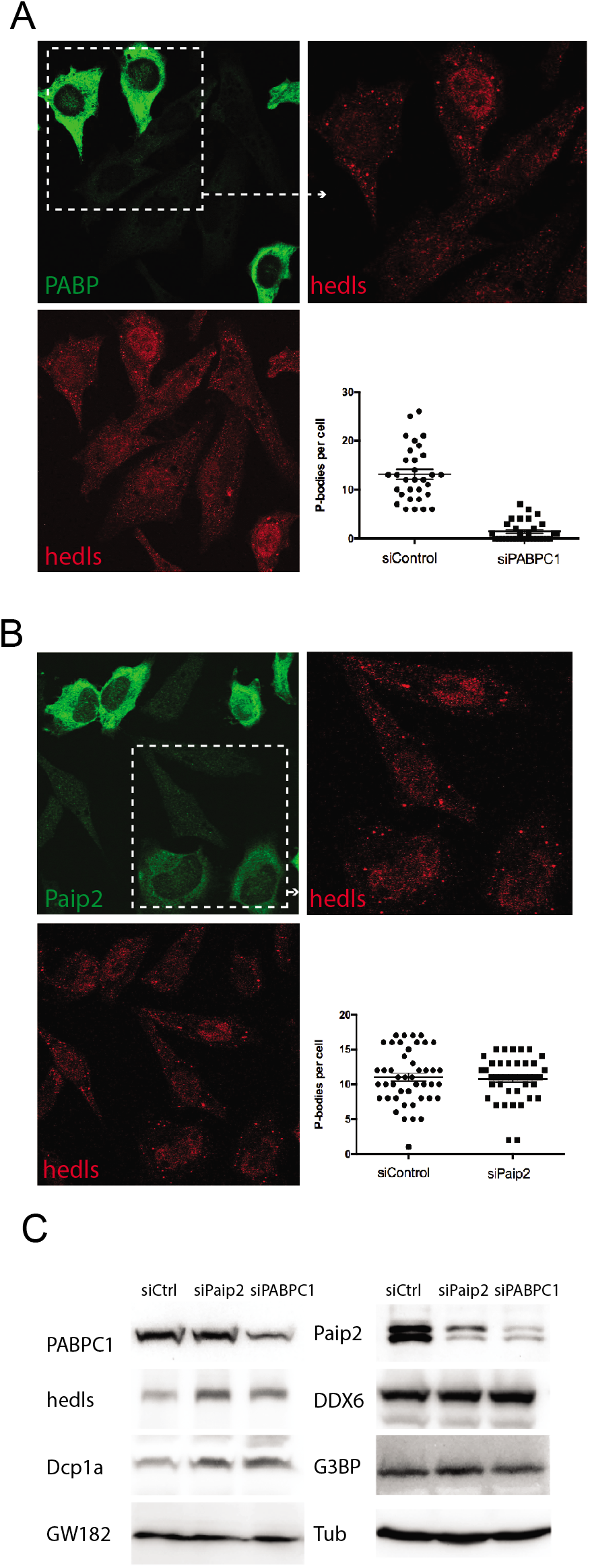
PB number and protein levels after PABPC1 or Paip2 depletion. (A-B) HeLa cells transfected with scramble siRNA were mixed with siPaip2 or siPABPC1 treated cells, to highlight knockdown effects and compare cells in the same environment. Staining of PABPC1 or Paip2 (*green*) indicated knockdown effects in cells. PB number was monitored by hedls *(red).* Visible P-bodies of 50 cells were counted manually and sample deviation was calculated. The PB number in siPABP cells was significantly lower at 0.01 significance level using two-tailed t-test for two samples with unequal variance. (C) Levels of PABPC1, Paip2, hedls, DDX6, Dcp1a, G3BP, GW182 and tubulin 72 hours after siRNA knockdown. The Paip2 antibody recognized both Paip2a and Paip2b.

### PBs cannot be re-induced in PABPC1 protein depleted cells

PB population may be reduced by siRNAs unrelated to their silencing activities and PBs can often be re-induced by stress (Serman et al., 2007). Therefore, we checked whether PBs could be re-induced upon cell stress. In mammalian cells, PBs and SGs can be clearly distinguished. PBs are relatively compact and dense, while SGs are bigger, loose and more irregular (Stoecklin and Kedersha, 2013, Souquere et al., 2009, Kedersha et al., 2005). When we treated PABPC1-depleted cells with arsenite to induce PBs and SGs, we were surprised to find that compact dense PBs failed to form. Instead, the PB component hedls was relocalized to SGs (Fig. 2). The colocalization was found across the cytoplasm by 3D confocal microscopy. This suggests that PB assembly is affected by depletion of PABPC1. To our knowledge, only DDX6, a RNA helicase also known as RCK/p54, can trigger PB/SG fusions and prevent PB formation when depleted (Mollet et al., 2008, Serman et al., 2007, Ayache et al., 2015). However, DDX6 protein levels remained unchanged after PABPC1 depletion (Fig. 1C). Thus, PABPC1 affects PB formation independently of DDX6.

**Figure 2.**
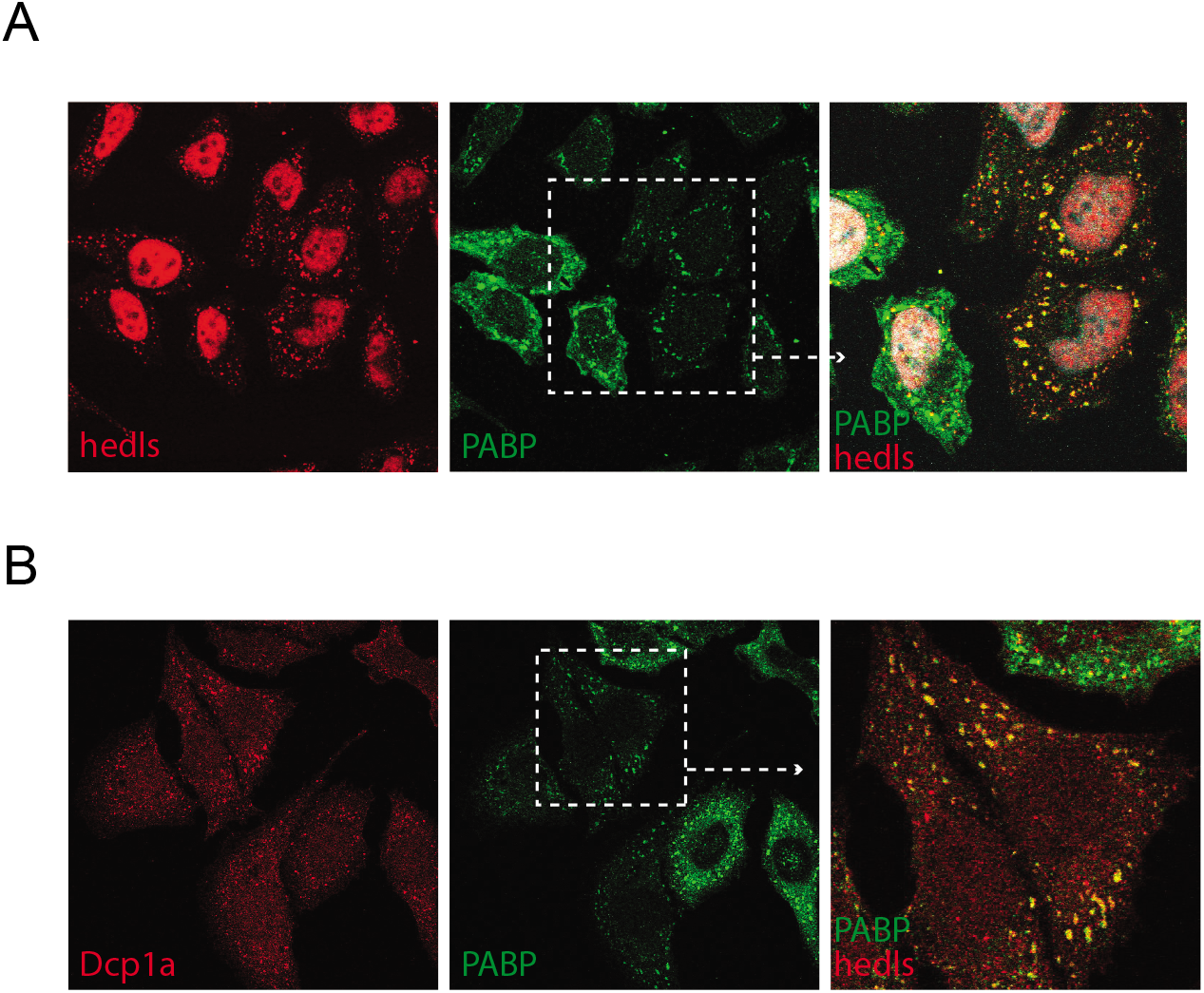
Relocalization of hedls or Dcp1a to stress granules in PABPC1-depleted cells. (A-B) HeLa cells were treated with 0.5 mM sodium arsenite for 0.5 hour before fixation. The PB components hedls (*red*) formed loose and irregular granules, colocalized to stress granules (*green*).

### SG assembly is not affected by PABPC1 depletion and is independent of PB assembly

Previously, it was observed that PABPC1 depletion induced SG in a low percentage of unstressed cells (Mokas et al., 2009). We checked SG assembly in stressed cells, and found SGs marked by G3BP, HuR (Fig. 3A & B) or Ago2 were not significantly affected by PABPC1 depletion. This implies that SGs can form in the presence of low levels of PABPC1 protein.

**Figure 3.**
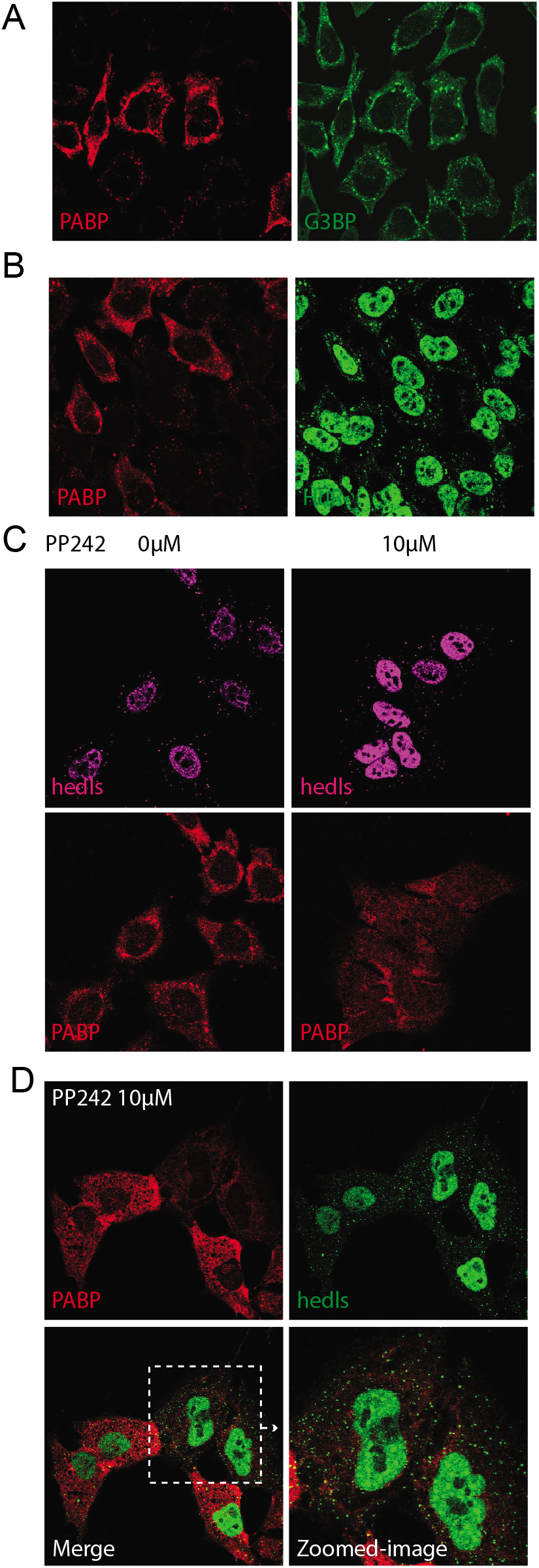
(A-B) Stress granule assembly is not affected by PABPC1 depletion. PABPC1 depleted cells (Red) displayed similar stress granule pattern to control cells, stained by anti-G3BP or anti-HuR (*green*). (C) P-bodies form independent of stress granules. HeLa cells treated by PP242 overnight failed to form stress granules *(red),* but had normal PBs present. (D) Separate PBs can be induced after SG inhibition in depletion of PABPC1. HeLa cells were treated with 10μM PP242 for overnight before stressed in 0.5 mM sodium arsenite for 0.5 hour. Individual PBs (*green*) formed in PABPC1 depleted cells (*red*).

The assembly of PBs and SGs are independent processes (Kedersha et al., 2005). PBs exist in unstressed cells in the absence of SGs. Meanwhile, PBs can be increased by arsenite stress in MEF cells expressing non-phosphorylated eIF2α mutant, which prevents SG formation (Kedersha et al., 2005). We treated HeLa cell with PP242, an mTOR inhibitor inhibiting SG assembly (Fournier et al., 2013, Feldman et al., 2009), and found PBs were increased by arsenite without SG formation (Fig. 3C). This implies that PBs do not require SGs to form in agreement with previous studies (Kedersha et al., 2005).

### Endogenous PABPC1 can be substituted by over-expressed PABPC1-GFP

We have shown that PB components can be found at PABPC1-rich SG (Fig. 2), and that only a fraction of the SGs contain PABPC1 in knocked-down cells (Fig. 3). Thus PBs require PABPC1, but not other SG components, for assembly.

It is known that overexpression of PABPC1 down-regulates the endogenous PABPC1 translation and protein abundance through self-regulation (Hornstein et al., 1999b, de Melo Neto et al., 1995, Wu and Bag, 1998, Hornstein et al., 1999a). This provides an opportunity to create cells where endogenous PABPC1 is replaced by exogenous PABPC1-GFP. We took advantage of an antibody (Abcam ab21060) that recognized the tail region of PABPC1 but not the chimeric PABPC1-GFP protein. Overexpression of PABPC1-GFP reduced endogenous PABPC1 to a very low level in cells (Fig. s1). However, the PBs assembled as normal, which implies that the over-expressed PABPC1-GFP can substitute endogenous PABPC1 in maintaining the PB population in unstressed cells.

### Separate PBs can be induced after SG inhibition

PBs assembly is closely related to PABPC1-containing SGs at low PABPC1 levels. We then asked whether structures of SG were required or not in such process. With PP242 inhibiting SG assembly, we stressed PABPC1-depleted cells with arsenite and individual PBs began to form (Fig. 3D). This shows that the low level of PABPC1 dissipated in cytoplasm can support PB assembly under stresses. It also confirms that SGs are not required for PB assembly. On the contrary, when the low level of PABPC1 was packed into SGs, it results in fusion of PBs and SGs. It is likely the tightly packed PABPC1 in SGs will not easily dissociate for mRNP remodeling, and thus lead to fusion of PBs and SGs.

### Overexpressed Dcp1a, TNRC6A and DDX6 can induce PB formation in PABPC1-depleted cells, and are found associated with the fusions of PBs and SGs

Overexpression of PB proteins can generate PBs that are not functional (Cougot et al., 2004). In PABPC1-depleted cells, overexpression of Dcp1a or DDX6 could induce PBs (Fig. 4A). When cells were stressed with arsenite, sorbitol, or heat shock, we found Dcp1a-GFP or HA-TNRC6A were present in PB/SG fusions (Fig. 4B & C). This confirms that multiple PB components were assembling at the fusion structures. Meanwhile, overexpression of Dcp1a or DDX6 could induce PB-like structures in stressed cells, as observed by other groups. Therefore, depletion of PABPC1 did not prevent the aggregation of overexpressed PB proteins into granules.

**Figure 4.**
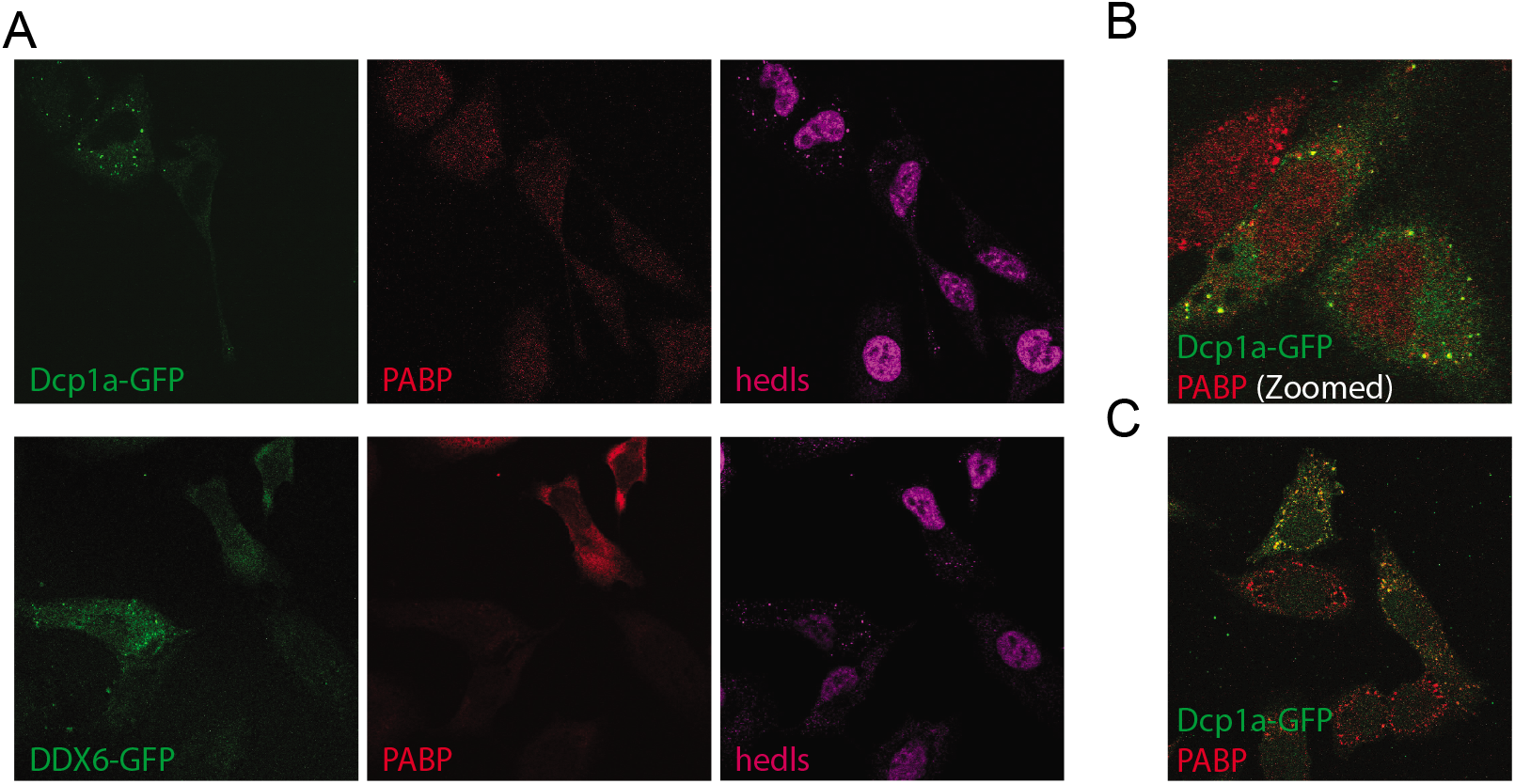
P-bodies can be induced in PABPC1 depleted cells. (A) Dcp1a or DDX6 overexpression (*green*) could induce P-body like granules in cells with low PABPC1 (*red*). (B-C) Dcp1a or TNRC6A (*green*) were partly localized to stress granules in arsenite stressed PABPC1 depleted cells. HeLa cells were transfected with siRNA and related plasmids simultaneously.

### Creating PB/SG fusions without PABPC1 depletion

We next asked how PABPC1, a protein absent from PBs, could affect their formation. One hypothesis is that PB assembly may initiate on PABPC1-containing mRNPs. When the residual PABPC1 after depletion is packed into SGs, PB components are recruited there and thus a PB/SG fusion emerges. We speculated that certain PB proteins interacting with PABPC1 might mediate the processes. We used a protein engineering approach to test our hypothesis.

GW182 proteins (TNRC6A/B/C in human) interact with PABPC1 through the MLLE domain and are required for PB assembly (Eulalio et al., 2007b; Liu et al., 2005a). The native affinity of the GW182 PAM2 motif for MLLE of PABPC1 is about 6 μM (Kozlov et al., 2010, Jinek et al., 2010). We engineered the PAM2 motif in GW182 to increase its affinity by including features from Paip2 PAM2 motif. The engineered GW182 protein contained a super-PAM2, binding PABPC1 about 300 times tighter (Fig. 5A, construct information in Materials and Methods). The super-PAM2 motif could efficiently localize fused GFP protein to stress granules (Fig. 5B). Introduction of super-PAM2 greatly increased association of GW182 and endogenous PABPC1 (Fig. 5C). When we stressed cells overexpressing super-GW182-GFP, the super-GW182 along with other PB components located to SGs to make PB/SG fusions (Fig. 6). This fusion is the combined result of the increased affinity PAM2/MLLE interaction and tightly packed PABPC1 at SG. It indicates that PABPC1 interacting proteins, like GW182, could play an important role in PB assembly through interaction with PABPC1.

**Figure 5.**
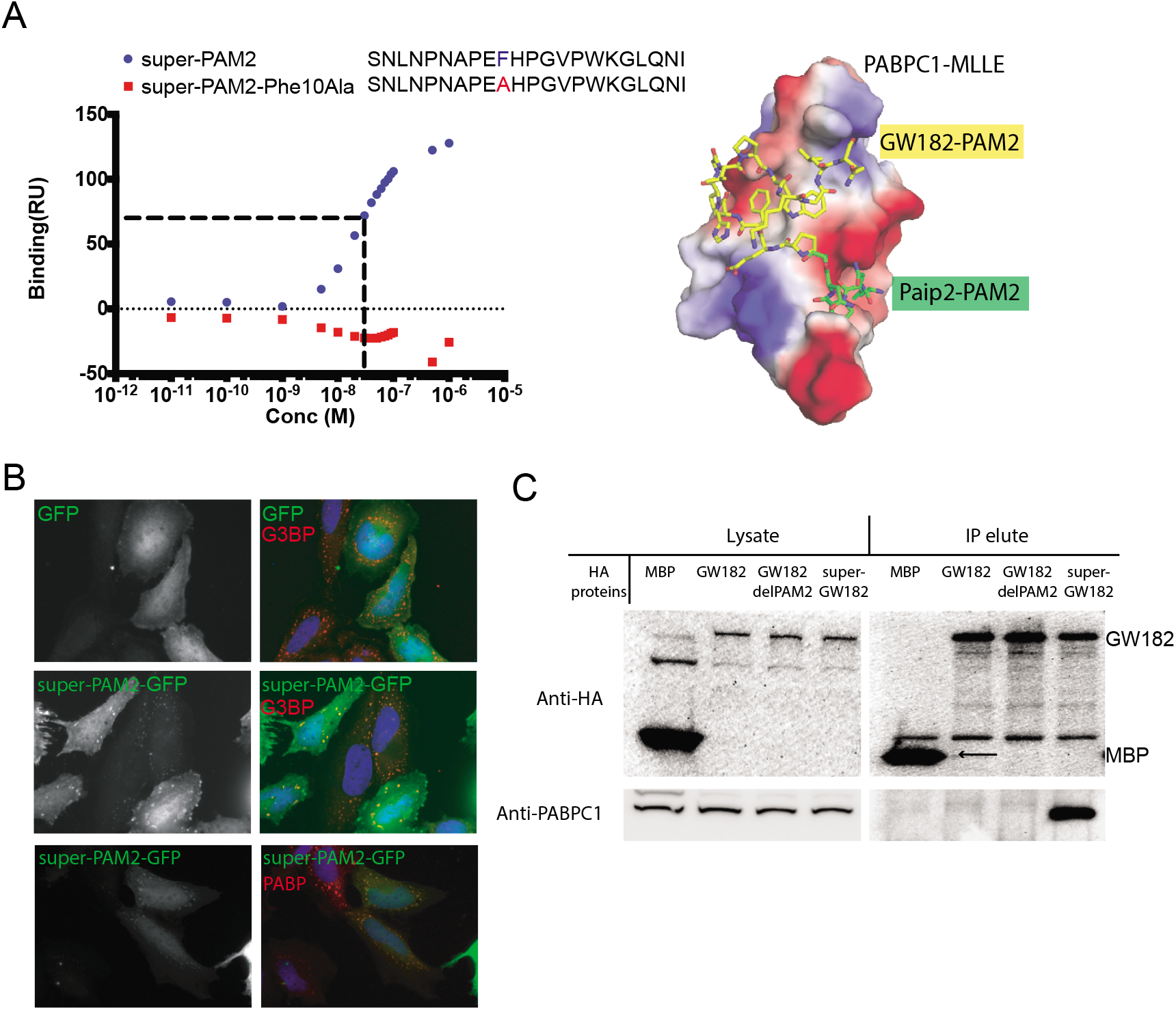
Engineering of super-GW182 to interact with PABPC1. (A) Super-PAM2 integrated sequences of Paip2-PAM2 and GW182-PAM2 to bind an extended surface on MLLE domain (PDB: 3KUS and 3KTP). Super-PAM2 bound MLLE of PABPC1 at around 20 nM measured by surface plasmon resonance using Biacore T100. The Phe10Ala mutant was not binding MLLE and used as control. (B) Overexpressed super-PAM2-GFP (*green*) localized to PABP or G3BP *(red)* marked stress granules upon sodium arsenite treatment. (C) HA-tagged MBP, GW182 (TNRC6A), GW182ΔPAM2 or super-GW182 were transfected in HeLa cells. Immunoprecipitation with anti-HA antibodies showed stronger interaction of super-GW182 with endogenous PABPC1. The weak association of natural GW182 and PABPC1 was not shown due to harsh washings.

**Figure 6.**
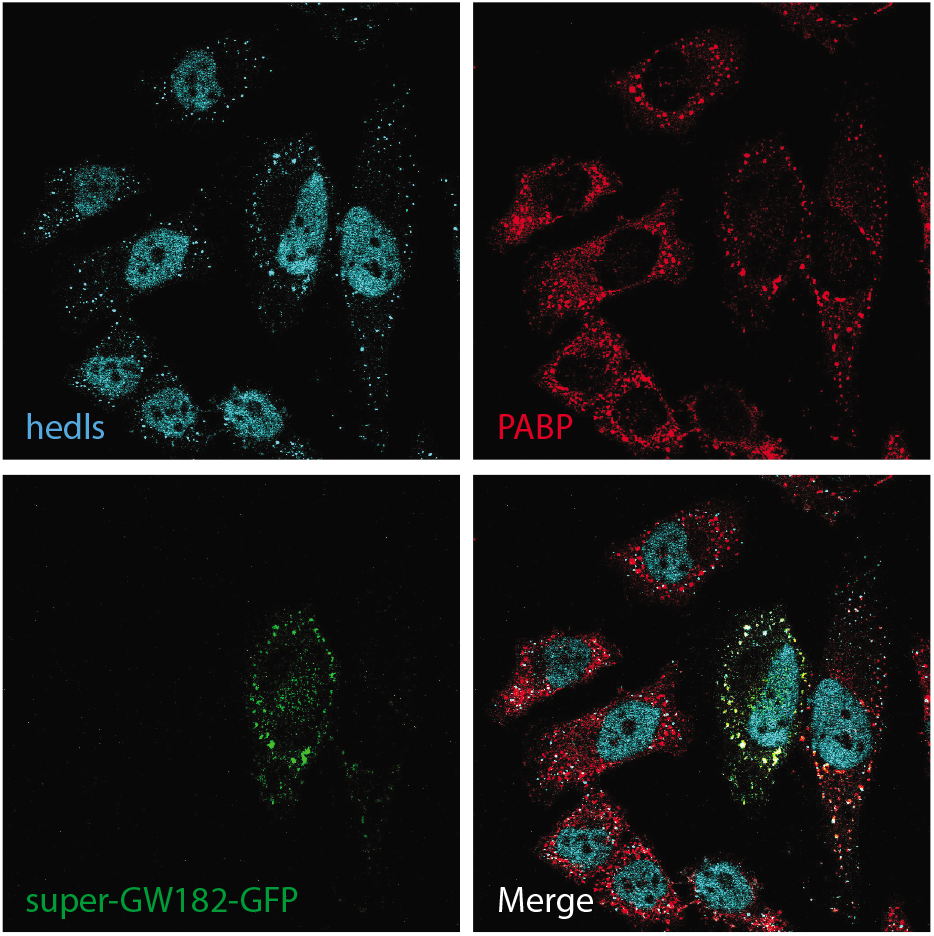
Simulation of P-body/Stress granule fusion using super-GW182. HeLa cells overexpressing super-GW182-GFP (*green*) were stressed by sodium arsenite and stained with anti-hedls (*cyan*) for P-bodies and anti-PABPC1 (*red*) for stress granules.

### Ago-binding region of super-GW182 is involved in recruiting other components

Both the N-terminal Ago-binding and mid Q/N region of GW182 were required for PB localization (Lazzaretti et al., 2009, Behm-Ansmant et al., 2006). We generated deletions of super-GW182 (TNRC6A) and checked their capability to bring PB/SG components together. Under arsenite stress, super-GW182ΔQ/N could induce fusion of PB/SG (Fig. 7B), while super-GW182ΔAgo was less efficient in fusing PB/SG (Fig. 7A). This suggests that the Ago-binding region is critical in subsequent PB components recruiting and assembly. This agrees with previous studies that showed that deletion of Q/N region in TNRC6A affects its PB localization much less than deletion of Ago-binding region (Lazzaretti et al., 2009). This indicates that the redirection of PB components to SGs by super-GW182 uses a similar mechanism to TNRC6A assembly into PBs.

**Figure 7.**
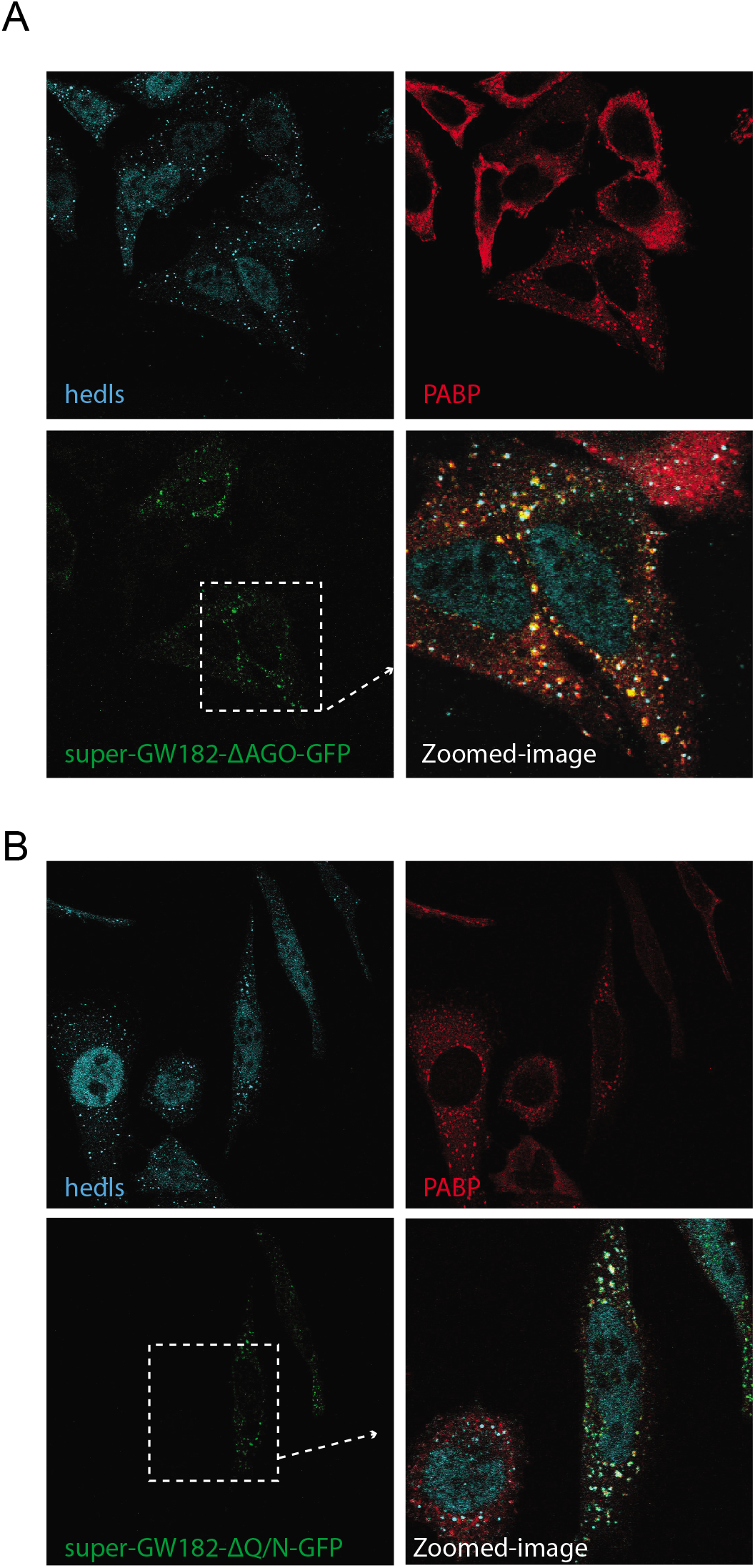
Ago-binding region of GW182 (TNRC6A) is important for recruitment of PB components. (A-B) Super-GW182ΔAgo or super-GW182ΔQ/N (*green*) was cotransfected with siPABPC1. PBs were labeled by anti-hedls (*cyan*) and stress granules were by anti-PABPC1 (*red*). Super-GW182ΔAgo was less efficient in redirecting PB components to SGs (A) compared with super-GW182ΔQ/N (B).

### GW182ΔPAM2 can aggregate in PABPC1 depleted cells and goes to SGs when stressed

We have shown that super-GW182 mimicked fusion of PB/SGs without depletion of PABPC1. This led us to ask what if the PAM2/MLLE interaction was missing? We transfected GW182ΔPAM2 (ΔT952–Q971) plasmid together with PABPC1 siRNA. HA-GW182ΔPAM2 formed PB-like granules in unstressed cells (Fig. s2A) and could be found in PB/SG fusions when stressed (Fig. s2B). The tendency of overexpressed PB proteins to aggregate into granules makes it hard to dissect the role of PAM2/MLLE interaction here.

## Discussion

Here, we have shown that PABPC1 protein level is critical to PB assembly. PABPC1 is a translation initiation factor and plays important role in mRNA metabolism. Depletion of PABPC1 decreases the number of PBs in unstressed cells. When the PABPC1-depleted cells are stressed, PB components are recruited to PABPC1-enriched SG. However, if SG assembly is inhibited, individual PBs can form in stressed cells.

PB assembly may initiate on PABPC1-containing mRNPs (Fig. 8). PABPC1 might be required in mRNP remodeling into P-bodies, thus PABPC1 depletion decreases PB numbers in unstressed conditions. The requirement of PABPC1 for PB formation drives PB components to certain SGs, where the residual PABPC1 is enriched after depletion. The displacement of PABPC1 at SG is difficult so individual PBs fail to form. When SG is inhibited by PP242, the same residual amount of PABPC1 in cytoplasm can support normal PB formation. It implies that the ability of PABPC1 to disassociate from mRNPs is required for formation of individual PBs. However, the mechanisms regarding how and when PABPC1 disassociates from mRNP are still to be investigated.

**Figure 8.**
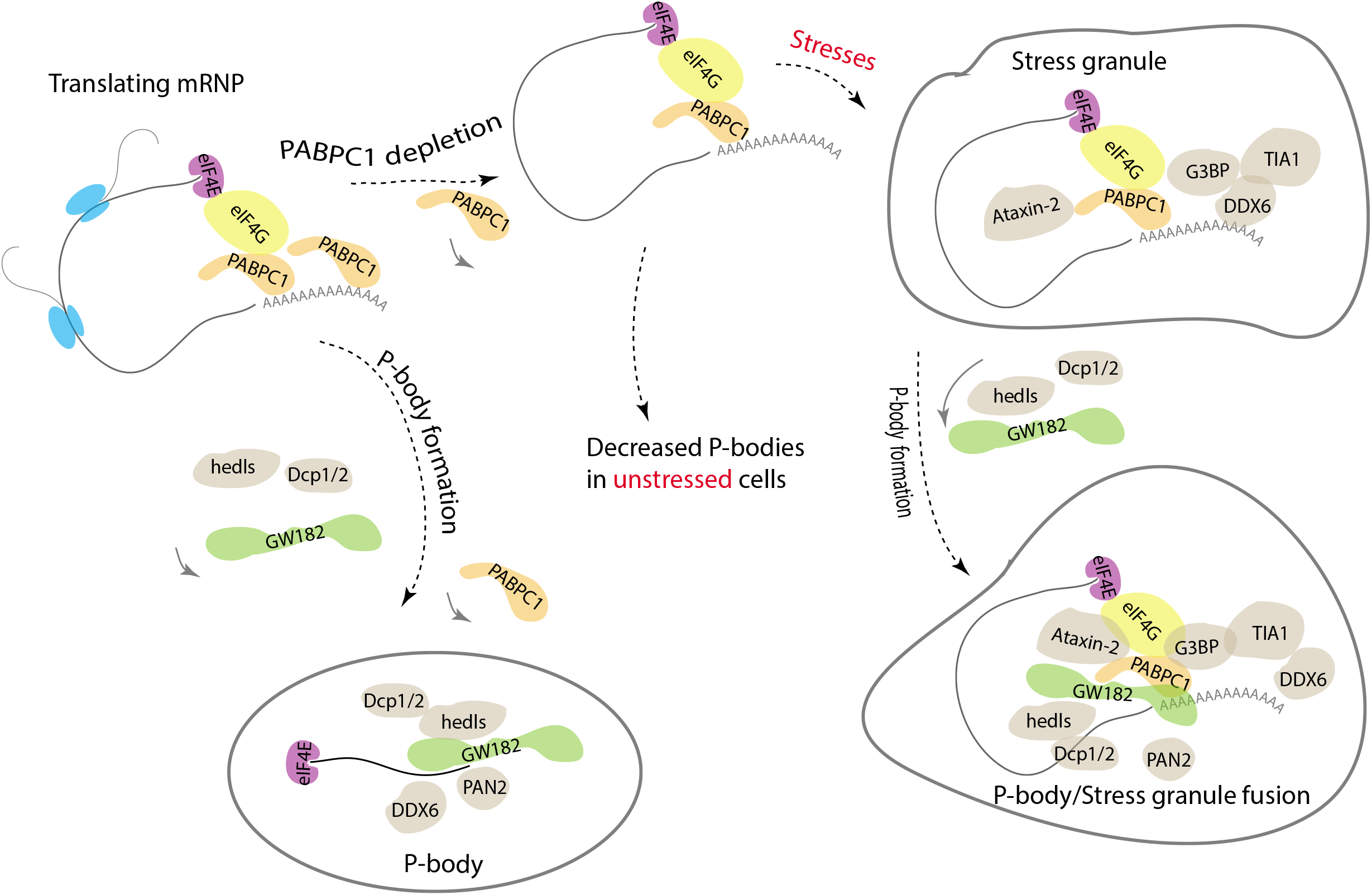
P-body formation on mRNPs or stress granules. PB assembly may initiate on PABPC1-containing mRNPs. The requirement of PABPC1 in PB formation drives PB components to stress granules, where the residual PABPC1 is enriched after depletion. However, the displacement of PABPC1 is difficult in SG and then individual PBs fail to form.

The displacement of PABPC1 from microscope-visible PBs is a critical step in PB assembly as was proposed by other colleagues (Chen and Shyu, 2013). PABPC1 plays a double role regarding poly(A) tail of mRNA. mRNA is stabilized and protected by PABPC1 from deadenylation by Ccr4/Not deadenylases (Tucker et al., 2002). Meanwhile, PABPC1 is required for deadenylation by Pan2/Pan3 deadenylases (Lowell et al., 1992, Zheng et al., 2008). The absence of PABPC1 in PBs is likely due to shortening of poly(A) in remodeling, as suggested by the absence of a signal for poly(A) RNA in PBs (Cougot et al., 2004).

Tendency of PB proteins to aggregate when overexpressed adds to the difficulty of PB assembly studies. Using super-PAM2 to relocate proteins to SGs is a novel way to evaluate their roles in PB assembly and dissect specific interactions. In all, we show that PABPC1 functions in mRNP remodeling and PB formation. Many mechanistic details need further studying. We look forward to future studies on roles of PABPC1 in mRNA metabolism.

## Materials and methods

### Plasmids and siRNA

pT7-EGFP-C1-HsTNRC6A, pT7-EGFP-C1-HsTNRC6B, pT7-EGFP-C1-HsRCK (DDX6), pT7-EGFP-C1-HsDcp1a, and pT7-EGFP-C1-HsDCP2 were gifts from Elisa Izaurralde (Addgene plasmid # 25030, 25031, 25033, 25034, 25035)(Tritschler et al., 2009). Sequence (agcaatctgaatccaaatgca) encoding amino acids SNLNPNA was inserted between nucleotides 4827A and 4828C of TNRC6A, or 4431T and 4432C of TNRC6B to create super-GW182. Reverse-PCR was used for constructing TNRC6AΔAgo (Δ7-300) and TNRC6AΔQ/N (Δ360-402). Plasmids expressing HA-MBP, HA-TNTC6A and HA-TNRC6AΔPAM2 were generous gifts from Drs. Eric Huntzinger and Elisa Izaurralde (Huntzinger et al., 2010). PABPC1 was inserted between BamH I and Not1 of pCDNA3-EGFP. siRNAs were synthesized at Dharmacon. The siRNA sequences used were siPABPC1 (5’-AAGGUGGUUUGUGAUGAAAAU-3’, 5’-AAUCGCUCCUGAACCAGAAUC-3’), siPaip2 (5’-GAGUACAUGUGGAUGGAAAUU-3’, 5’-UGGAAGAUCUUGUGGUCAAUU-3’). Control siRNA was purchased from Qiagen (SI03650318).

### Cell culture and transfections

HeLa S3 or MEF cells were cultured in DMEM supplemented with 10% fetal bovine serum. 10^5^ cells were plated per well in 24-well plate the day before transfection. 60-90 pmol siRNA or 0.8 μg DNA plasmid was mixed with 2 μl Lipofectamine 2000 in Opti-MEM and then added to cells. After 48 hours, cells were trypsin digested and split onto cover slides. Cells were treated and fixed after 24 hours. For western analysis, cells were changed into fresh medium for another day and harvested in SDS loading buffer.

### Immunofluorescence and confocal microscopy

Cells were fixed with cold methanol (−20°C) for 10 minutes and then treated with 0.1% Triton X-100 in 1x PBS. Cells were blocked with 5% normal goat serum (Millipore S26) in PBS. Cells were incubated in 1x PBS, supplemented with anti-PABPC1 (Abcam ab21060; Santa Cruz sc32318)(1:200), anti-hedls (Santa Cruz sc8418)(1:1000), anti-Paip2 (Sigma-Aldrich P0087)(1:500), anti-eIF4G1 (Sigma-Aldrich AB-1232)(1:200), anti-Ataxin-2 (BD Biosciences 611378)(1:200), anti-HA (Covance MMS-101P)(1:200), anti-Dcp1a (Abcam ab47811)(1:200), anti-DDX6 (Abcam ab40684)(1:200), anti-EDC4 (hedls)(Abcam ab72408)(1:200), anti-Ago1 (Santa Cruz sc53521)(1:200), anti-G3BP (gift of Dr. Imed Gallouzi)(1:500) or anti-HuR (gift of Dr. Imed Gallouzi)(1:200). Cells were washed in PBS 3 times, before incubated with corresponding second antibodies conjugated with Alexa488 or Alexa647 (Sigma-Aldrich A31620, A31628, A31571) or Dylight550 (Bethyl A120-101D3) or Rhodamine (Millipore 12-509, 12-510) at 1:200 – 1:500 dilutions. DAPI (Roche) was added to washing buffer at 0.5μg/ml to treat cells for 10 minutes. Cover slides were finally mounted in ProLong Gold antifade reagent (Life technology P36930). Images were collected on Zeiss LSM 310 confocal microscope in the McGill University Life Sciences Complex Advanced BioImaging Facility (ABIF). PBs were counted manually in a sample size of 50 cells for each group. Sample deviation was calculated within group.

### Western blotting

Protein samples were heated at 95°C and separated in SDS-PAGE. Proteins were then transferred to PVDF membrane in Tris/glycine buffer, with 20% methanol in cold room. PVDF membrane was blocked in 1x TBS (pH 7.5), containing 0.05% Tween-20 and 5% skim milk powder or bovine serum albumin. Besides antibodies above, anti-GW182 (Novus NBP1-88232)(1:200) and anti-tubulin (Sigma-Aldrich T9028) were also used for detection of related proteins.

### Immunoprecipitation

Cells were lysed in 20 mM Hepes (pH 7.4), 150 mM NaCl, 0.5% NP-40, 2 mM DTT, 2 mM MgCl_2_, 1 mM CaCl_2_ and protease inhibitor tablet (Roche), and cleared by centrifugation. Optimized amount of antibodies were added to cleared lysate for 2 hours. Dynabeads protein A or protein G were washed and added to lysate for 0.5 hour. Dynabeads were then washed with 1x PBS and boiled in SDS loading buffer for further analysis.

### Surface plasmon resonance

GST-MLLE protein was applied to Series S sensor chip CM5 bound with anti-GST antibodies (GE Healthcare BR100223), in Biacore T100. After wash, various concentrations of super-PAM2 (SNLNPNAPEFHPGVPWKGLQ) or super-PAM2-Phe10Ala (SNLNPNAPEAHPGVPWKGLQ) were added to bind GST-MLLE captured on chip. The corresponding steady states measured in relative units (RU) were plotted versus concentrations of peptides in the flow system to estimate dissociate constants.

## Acknowledgements

We acknowledge Drs. Elisa Izaurralde, Witold Filipowicz and Imed Gallouzi for generously sharing plasmids or antibodies. We are grateful for suggestions and help from colleagues during the project.

## Author contributions

J.X. designed the experiments. Y.C. and X.W. assisted J.X. in carrying out the experiments. G.K. designed the super-PAM2 sequence. J.X. and K.G. prepared the manuscript.

