## Supplemental figures for "Poly(A) binding protein is required for mRNP remodeling to form P-bodies in mammalian cells"

**Figure legends of supplementary figures**

Figure s1. Lowered endogenous PABPC1 by overexpressed PABPC1-GFP does not affect PB formation. Anti-PABPC1 (Abcam ab21060) revealed low level of PABPC1 (*red*). PBs were shown by anti-hedls (*magenta*).

Figure s2. Overexpressed GW182ΔPAM2 can induce P-body like granules in unstressed cells (A) and be found at stress granules in depletion of PABPC1 (B). HeLa cells were co-transfected with GW182ΔPAM2 (*green*) and control siRNA or siPABPC1. Cells were stained with anti-PABPC1 (*red*) to indicate protein level and stress granules.

### Figure s1

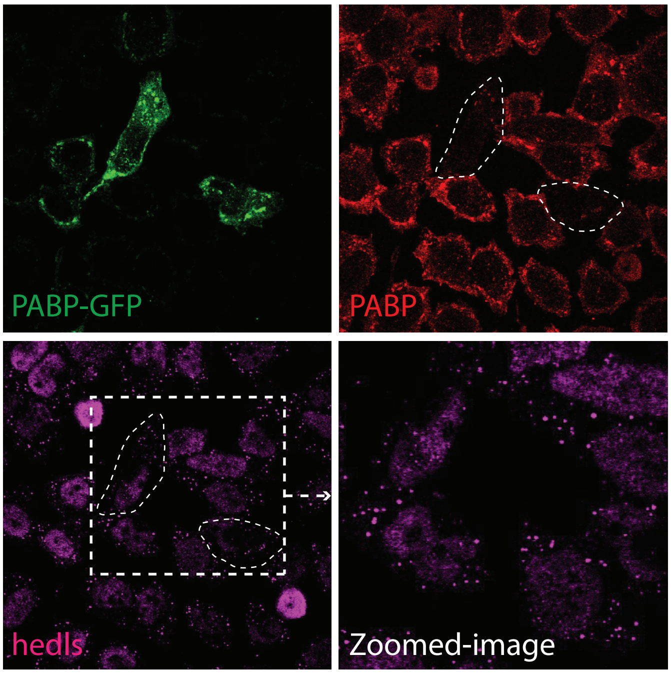

Figure s2

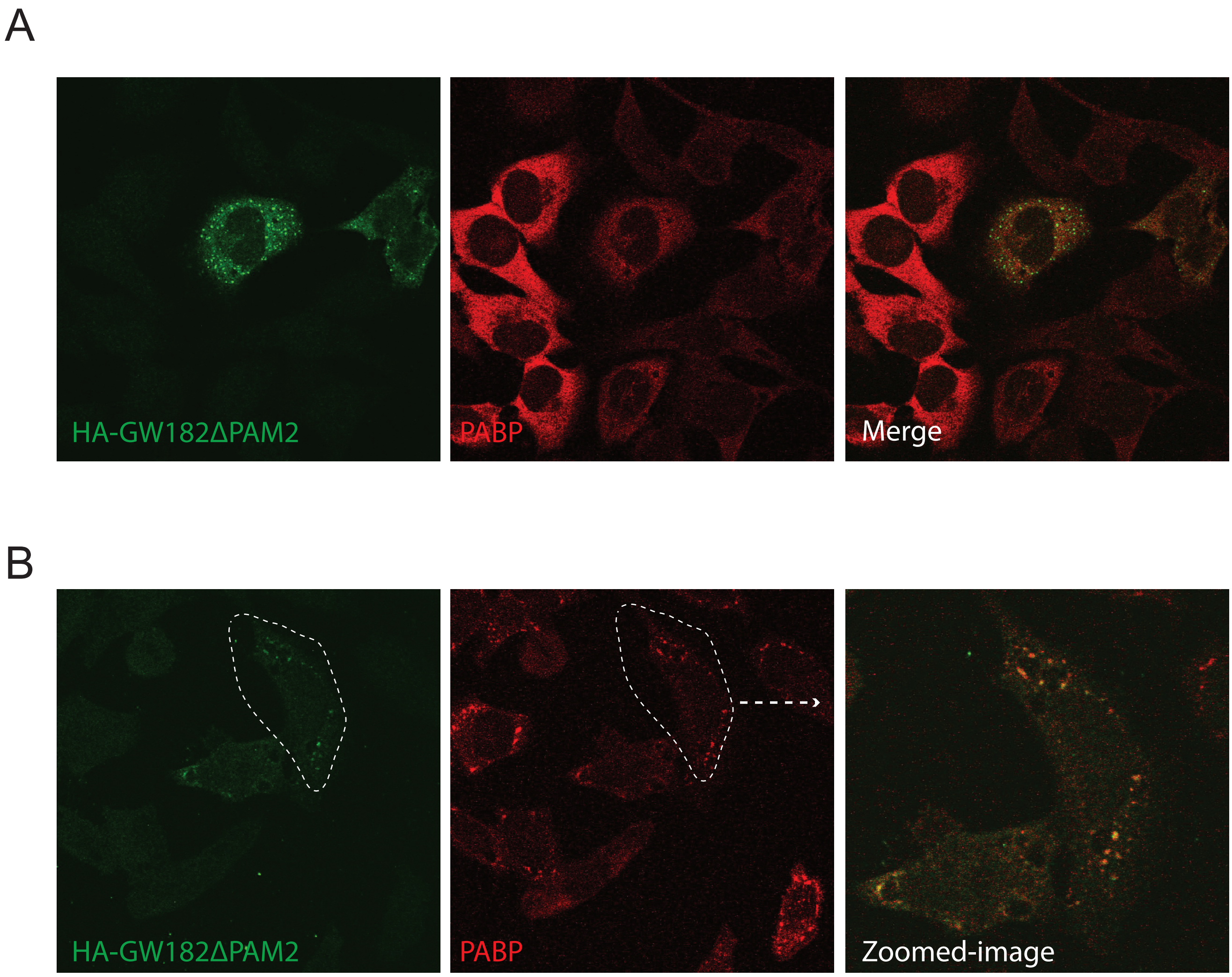
A
HA-GW182ΔPAM2
PABP
Merge
B
HA-GW182ΔPAM2
PABP
Zoomed-image
